# Investigating regions of shared genetic variation in attention deficit/hyperactivity disorder and major depressive disorder: A GWAS meta-analysis

**DOI:** 10.1101/2020.04.22.054908

**Authors:** Victoria Powell, Joanna Martin, Anita Thapar, Frances Rice, Richard J. L. Anney

**Author notes:** Corresponding Author: Victoria Powell (UK).

## Abstract

Attention deficit/hyperactivity disorder (ADHD) demonstrates a high level of comorbidity with major depressive disorder (MDD). One possible contributor to this is that the two disorders show high genetic correlation. However, the specific regions of the genome that may be responsible for this overlap are unclear. To identify variants associated with both ADHD and MDD, we performed a meta-analysis of GWAS of ADHD and MDD. All genome wide significant (p=5×10^−8^) SNPs in the meta-analysis that were also strongly associated (p=5×10^−4^) independently with each disorder were followed up. These putatively pleiotropic SNPs were tested for additional associations across a broad range of phenotypes. Fourteen linkage disequilibrium-independent SNPs were identified that were associated with each disorder separately (p=5×10^−4^) and in the cross-disorder meta-analysis (p=5×10^−8^). Nine of these SNPs had not been reported previously in either individual GWAS and can be considered as novel signals. Evidence supported nine of the fourteen SNPs acting as eQTL and two of the SNPs as brain eQTL. Index SNPs and their genomic regions demonstrated associations with other mental health phenotypes. Through conducting meta-analysis on ADHD and MDD only, our results build upon the previously observed genetic correlation between ADHD and MDD and reveal novel regions of the genome that may be implicated in this overlap.

## Introduction

DSM-5 defined neurodevelopmental disorders that typically onset early in development demonstrate a high level of comorbidity with later-onset psychiatric disorders (1). Attention deficit/hyperactivity disorder (ADHD) is one such neurodevelopmental disorder and is associated with multiple long-term poor health outcomes, including depression (2). Depression is a leading cause of disability worldwide (3) and its incidence rapidly increases during adolescence and peaks in early adulthood (4,5). Evidence of the association between ADHD in childhood or adolescence and increased risk of subsequent depression has been found in both general population (6) and clinical samples (2,7–9). Moreover, clinical outcomes for those with comorbid ADHD and depression are often worse than in individuals with depression alone (7,10). The factors that underlie this strong association between ADHD and depression are unclear.

As ADHD and depression are both heritable and familial (11–14), one contributing factor that could partly explain the relationship between ADHD and depression is shared genetic risk. Early evidence for this came from twin studies, which observed a genetic correlation ranging between 0.23 to 0.77 for ADHD and depression diagnoses or symptoms in children and adolescents (15–19). More recently, genome-wide association studies (GWAS) have supported the genetic overlap of ADHD and major depressive disorder (MDD; used synonymously with ‘depression’ in the current study), with an estimated ADHD-MDD genetic correlation of r_g_=0.42 (SE=0.03) (13,14). Moreover, studies have shown that polygenic scores derived from the ADHD GWAS are associated with depression diagnosis in an adult population sample (20), depression symptoms in an adolescent population sample (21) and depression symptoms in a twin sample of children (22). In line with these findings, the most recent GWAS meta-analysis of eight psychiatric disorders by the Cross Disorder Group of the Psychiatric Genomics Consortium (PGC) (23) reported a similar robust genetic correlation between ADHD and depression (r_g_=0.44, SE=0.03). Given the focus by Lee et al., (2019) on exploring cross-disorder effects including eight psychiatric disorders and reporting based on statistics across all eight diagnoses, regions impacting only ADHD and MDD are likely to be omitted from the key findings due to insufficient evidence across the remaining six diagnoses. Risk mechanisms that explain the link between ADHD and depression might also involve other phenotypes that have not been previously studied; for example, psychiatric phenotypes such as early irritability and anxiety are known antecedents of depression, including in those with ADHD (24–26).

In this paper we conducted a GWAS meta-analysis of ADHD and MDD, aiming to identify novel associated SNPs with contribution to both disorders. Our secondary aim was to investigate associations between SNPs implicated by this cross-disorder meta-analysis and a range of additional GWAS phenotypes.

## Methods

### Samples and Measures

We used published GWAS summary statistics of ADHD (14) and MDD (13). GWAS data were downloaded from https://www.med.unc.edu/pgc/results-and-downloads/. Additional local quality control of the downloaded data was performed to align genome strand, genome build (hg19) and marker nomenclature (HRC1.1) across the studies. All analysis was limited to autosomal SNPs.

#### Attention deficit/hyperactivity disorder: Demontis et al., 2019 (14)

Demontis and colleagues identified 12 genome-wide significant loci associated with ADHD by combining 12 cohorts (20,183 cases and 35,191 controls), mainly of European and North American ancestry and one of Chinese ancestry. The Danish population-based cohort (iPSYCH) used health records from the Danish Psychiatric Central Research Register to identify those individuals diagnosed with ICD-10 ADHD. The remaining 11 cohorts consisted of 4 parent-offspring trio cohorts and 7 case-control cohorts in which individuals with ADHD were recruited from clinics, hospitals, or medical registers, and were diagnosed using standard tools administered by trained researchers or clinicians. Imputation of non-genotyped markers was conducted using the 1000 Genomes Project Phase 3 reference panel. Heterogenous ancestry in samples has been shown to give rise to population stratification bias in genome-wide analyses (27). Thus, we restricted our analysis to samples of European ancestry only, leaving 19,099 cases and 34,194 controls.

#### Major depressive disorder: Wray et al., 2018 (13)

Wray and colleagues identified 44 genome-wide significant loci associated with MDD by combining 7 case-control cohorts (135,458 cases and 344,901 controls). The first cohort was a mega-analysis of 29 European ancestry samples where MDD cases were required to meet DSM-IV, 1CD-9 or ICD-10 criteria using structured diagnostic interviews, clinician-administered checklists or a review of medical records. The 6 additional cohorts were also of European ancestry where cases were defined according to DSM-IV, ICD-9 or ICD-10 diagnoses of MDD derived via interviews, self-report, receiving medical treatment and national or hospital treatment records. The published data includes an MDD cohort from the 23andMe study. Due to data restrictions, we excluded the 23andMe cohort from our analysis, leaving a sample of 59,851 cases and 113,154 controls.

### Analysis

#### SNP-based heritability and genetic correlation

SNP-based heritability (SNP h^2^) and the genetic correlation of ADHD and MDD GWAS were calculated using the LD-score approach (28). Estimates were calculated on the liability scale using a population prevalence of 0.05 for ADHD and 0.15 for MDD. Summary data were harmonised to common build, strand and nomenclature and SNP h^2^ estimates calculated limited to the HapMap-3 SNP subset provided by the LDSC software.

#### GWAS meta-analysis to identify regions of joint association

Our aim was to identify regions of common association between ADHD and MDD, regardless of the direction of the effect. Strong associations in opposing directions are of biological interest and may also be expected where one trait could act as a counter measure of the second (e.g. wellbeing and depressive symptoms). Therefore, we meta-analysed the maximum SNP effect in a GWAS summary-data-based meta-analysis. We performed the meta-analysis using a fixed-effects inverse variance-weighted model using METAL (29) with ADHD used as the index. A fixed-effects model rather than a random-effects model was chosen to maximise power to detect SNPs associated with ADHD and MDD (30). A genome-wide significance (GWS) threshold of 5×10^−8^ was used.

For each of the identified SNPs from the MDD-ADHD meta-analysis, we compared the association findings against the original ADHD and MDD GWAS. Based on these comparisons we describe three classes of association. Model 1 is defined as a GWS meta-analysis hit where the p-value for ADHD GWAS is smaller than both the MDD GWAS and meta-analysis p-values, thus appearing to be more robustly implicated in ADHD than the combination of both phenotypes. Model 2 is alike to Model 1, but with the MDD p-value as the smallest. Model 3 is defined as a GWS meta-analysis hit where the meta-analysis p-value is smaller than both the ADHD and MDD GWAS p-values. Model 3 is suggestive of a variant contributing to both phenotypes and highlights potential regions of joint association between ADHD and MDD. Thus, this is our model of interest and we report all SNPs meeting Model 3 criteria. To limit the identification of putatively stochastic Model 3 associations, we limited reporting to SNPs that show an ADHD and MDD maximum p-value of 5×10^−4^.

SNPs that met Model 3 criteria were Linkage Disequilibrium (LD) pruned using PLINK (version 1.07 (31)). LD statistics for pruning were derived from the European super population 1000 genomes project phase 3 reference genotypes. “LD ranges” were defined as regions around the index association containing SNPs with r^2^≥0.2 with the primary SNP and p<.05.

Due to concerns in the literature regarding over-reliance on p-values (32), we repeated our meta-analysis using z-score thresholds to define Model 1, 2 and 3 SNPs (Supplementary File 1: Table S1).

#### Annotation of association signals within identified regions

To aid in the biological interpretation of the associated markers, the index SNPs, SNPs in high LD with the index SNP, and LD ranges were annotated with data related to both physical and functional landmarks. SNPs in high LD were defined as those SNPs with r^2^ >0.8 with the index SNP within the European super population 1000 genomes project phase 3 reference genotypes. Gene transcript, cis-eQTL, variant consequence and chromatin state annotations were investigated as described in Supplementary File 2.

#### GWAS association

Regions of GWS association in the meta-analysis (LD ranges containing index association of p<5×10^−8^) were mapped to strong associations (LD ranges containing index association of p<1×10^−5^) for a set of 37 reference GWAS of human phenotypes. These GWAS are heritable, reasonably well-powered, complex human traits that could be of relevance to both ADHD and MDD. The studies covered 5 categories of traits: mental health (13 traits), personality (5 traits), cognitive (3 traits), anthropometric (4 traits) and ‘other’ (12 traits). The ‘other’ category included traits relating to physical activity, smoking and health conditions of the skin, digestive, genitourinary, metabolic, immune and nervous systems. All GWAS LD ranges were calculated as described previously and overlaps were mapped using bedtools (https://bedtools.readthedocs.io/). Moreover, we directly assessed the association at the index SNPs against all 37 GWAS. All SNPs were p-corrected < 0.05. Bonferroni correction based on 37 independent tests was applied (Supplementary File 3). A full list of the GWAS used in this analysis is included in Supplementary file 1 (Table S2).

## Results

### SNP Heritability and Genetic Correlation

After quality control of the ADHD summary data, 5,907,045 SNPs remained. ADHD SNP h^2^ measured on the liability scale was estimated as h^2^=0.22 (0.02). After quality control of the MDD summary data, 7,104,680 SNPs remained. MDD SNP h^2^ on the liability scale was h^2^=0.15 (0.01). Genetic correlation between ADHD and MDD was estimated as r_g_=0.52 (0.04).

### Identification of Regions Showing Evidence of Common Association

We identified 14 SNPs that met Model 3 criteria in the combined analysis (sample size per SNP range of 120,401 to 191,525); that is they were associated at GWS level in the combined analysis, the p-value was lower in the combined analysis compared to the individual GWAS, with p<5×10^−4^ for ADHD and MDD separately (see Table 1). Each of the 14 SNPs demonstrated concordant direction of effect for ADHD and MDD. Nine of the 14 SNPs were novel in that they were not previously identified as being GWS or within regions that harbour GWS association for ADHD or MDD (see Supplementary Table S3). For each of the 14 index SNPs, the regions that contained them were associated at the GWS level in at least one of the 37 reference GWAS, in addition to ADHD and MDD (see Table S4). Moreover, these associations were also observed for the same SNP (see Supplementary File 3). We performed follow-up analyses for each of these 14 index SNPs, which we describe below and in the Supplementary Materials. For 2 of the SNPs which were of greatest interest – the strongest (smallest p-value) meta-analysis signal (rs12658032) and the strongest signal found to act as a brain eQTL (rs8084351) – we also present detailed figures of key results. Detailed results for all other SNPs are in the Supplementary Materials.

**Table 1.**
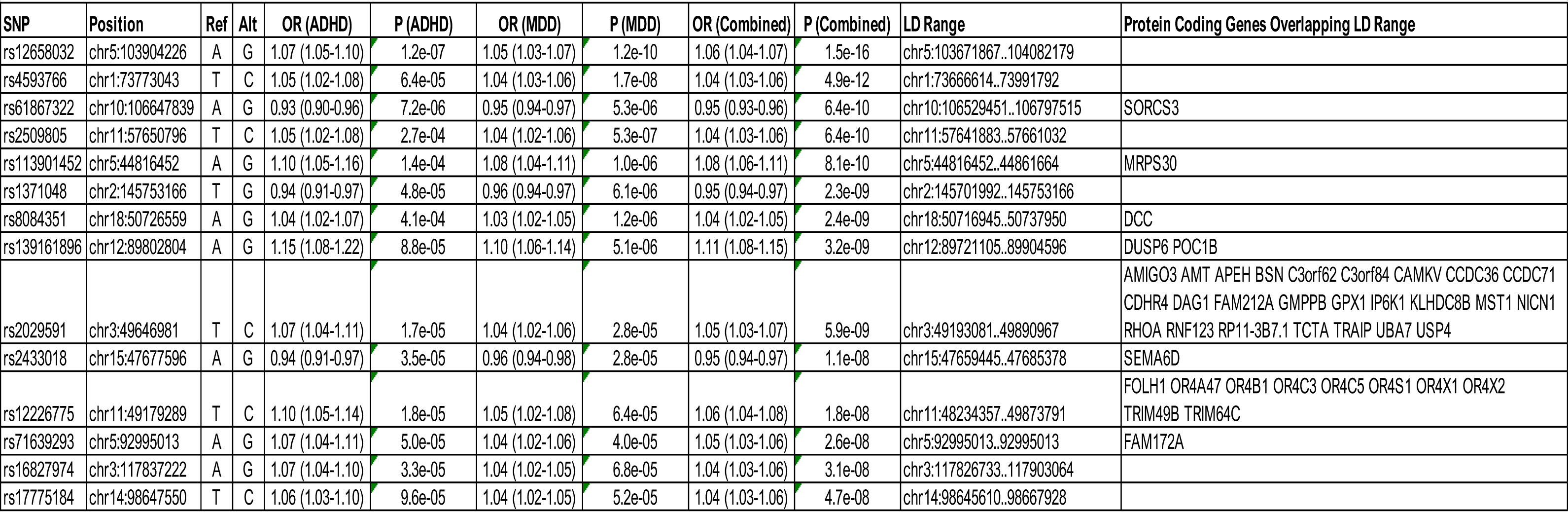
Summary of GWS associations for SNPs identified as contributing to both ADHD and MDD. Protein coding genes located in the immediate region of each SNP are reported.

Two of these 14 index SNPs (rs12658032 and rs4593766) had previously shown strong evidence for association with the individual ADHD or MDD GWAS. The first of these SNPs, rs12658032, (OR_meta_=1.06(1.04-1.07); P_meta_=1.54×10^−16^) was the strongest association observed, with GWS association observed in MDD and strong association for ADHD (see Table 1; Figure 1). The SNP is mapped to chromosome 5q21.2 in a region showing no protein-coding annotation, no evidence of eQTL (P<10^−5^) (Supplementary file 1: Tables S5-7) and no evidence of open chromatin (Table S8). The region shows overlap with the lincRNA transcripts *RP11-6N13.1* (ENSG00000251574.6) and *RP11-6N13.4* (ENSG00000253776.1) (Table S9-10). Based on data from the GTEx Portal, *RP11-6N13.1* and *RP11-6N13.4* are expressed predominantly in the testis. There is no evidence to support expression in the brain. rs12658032 shows SNP-level associations in additional mental health-related GWAS in the same direction of effect as observed in ADHD and MDD GWAS, including depressive symptoms and bipolar disorder, as well as anorexia in the opposing direction of effect (see Supplementary File 3; Figure 2). The region is also associated with years of education, birthweight, BMI and waist circumference (Supplementary file 1: Table S4).

**Figure 1.**
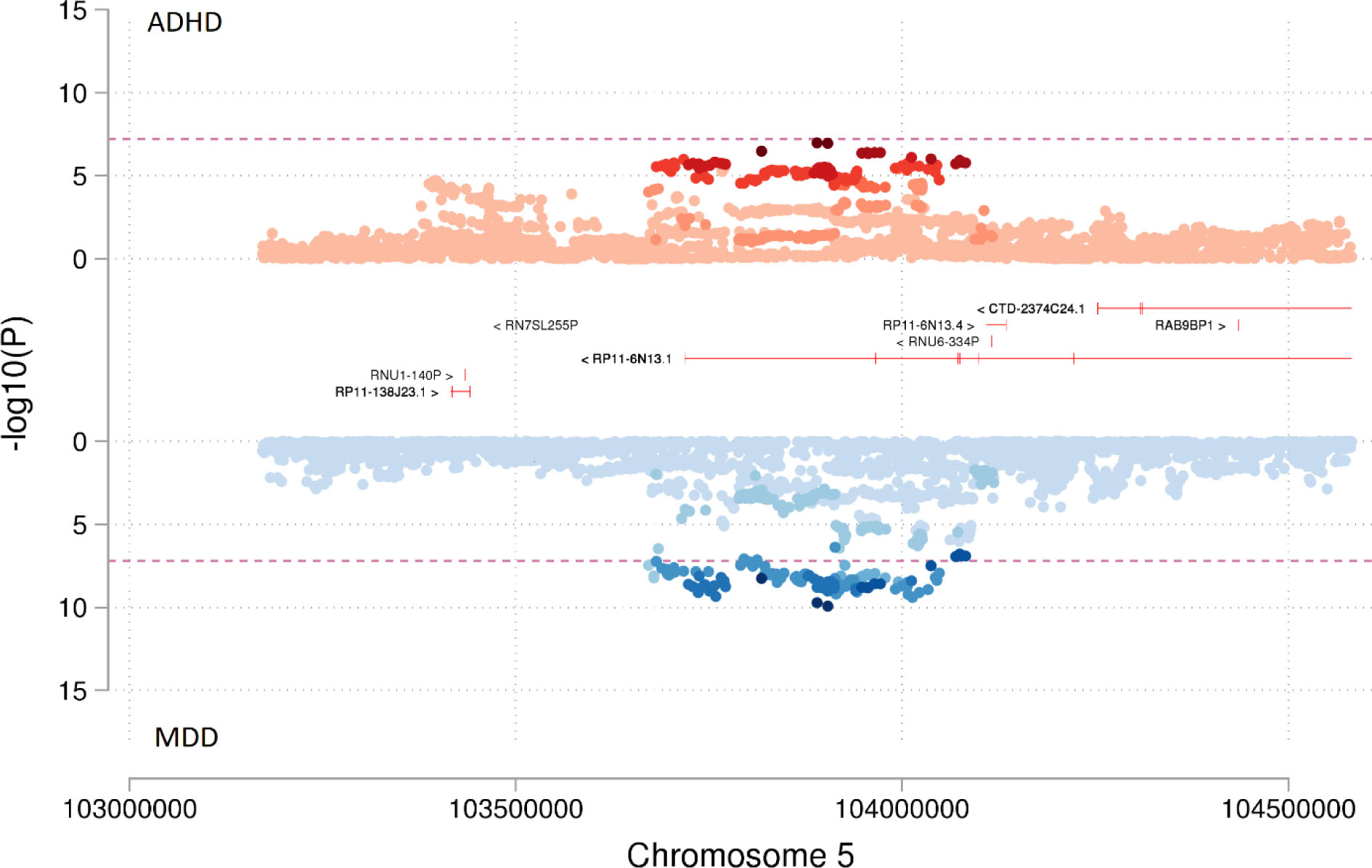
Miami plot showing the association of SNP rs12658032 with ADHD compared to MDD. Miami plot from the meta-analysis of ADHD and MDD GWAS. SNP rs12658032 on chromosome 5 was associated with both ADHD (p=1.15×10^−7^) as shown in red and with MDD (p=1.18×10^−10^) as shown in blue to a genome-wide significant level and was also genome-wide significant in meta-analysis (p=1.54×10^−16^). This SNP was located in a protein-coding gene desert. Non-protein-coding genes are shown in red. Genes are defined according to residence within the LD block of index SNP. The x-axis is chromosomal position in base pairs and the y-axis is the p value (-log10 p value) of the association of the SNP with both disorders.

**Figure 2.**
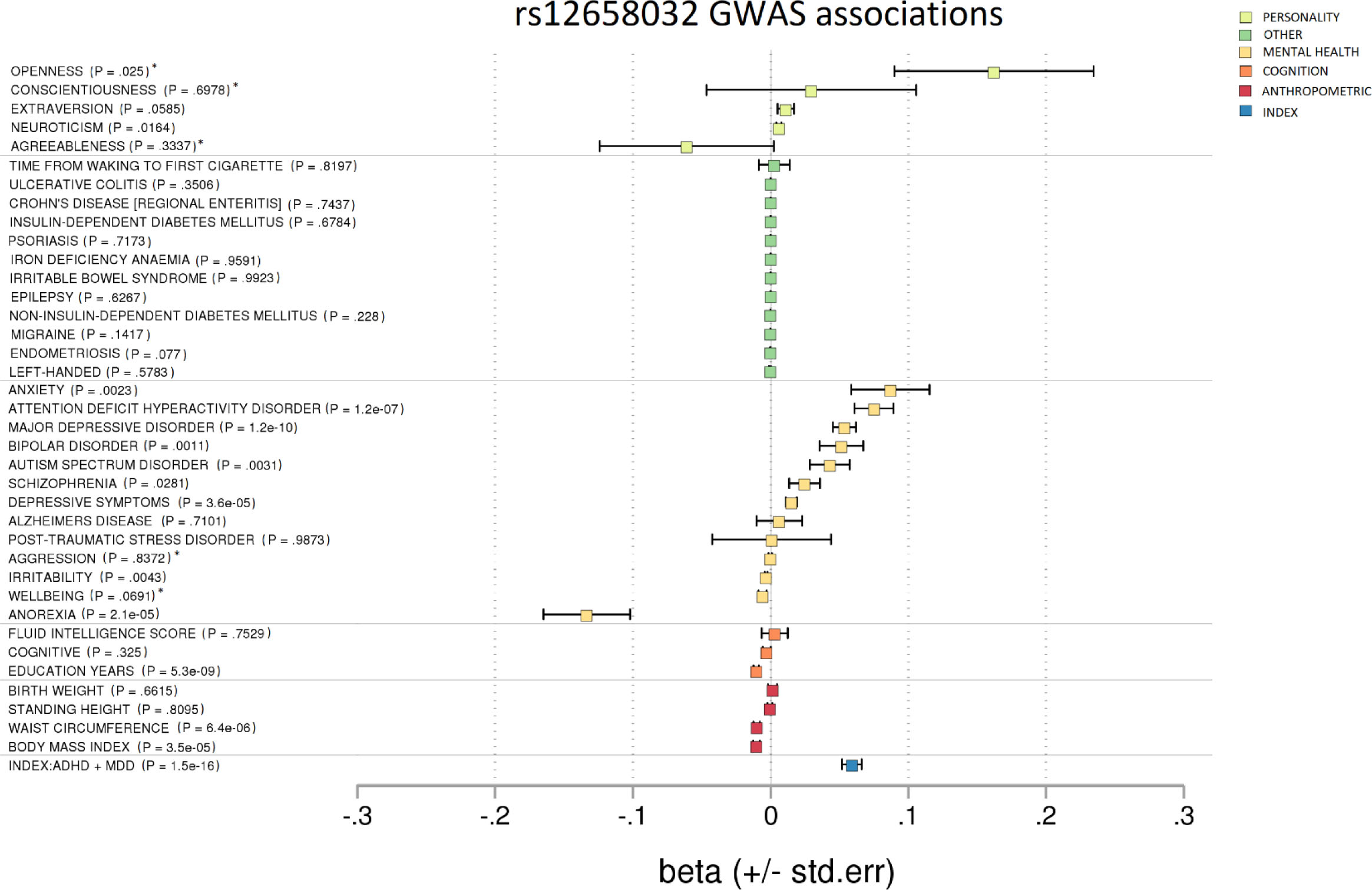
Forest plot showing GWAS associations of rs12658032. To investigate pleiotropy, rs12658032 was tested for associations across 37 phenotypes covering various aspects of health and functioning. rs12658032 was associated mainly with mental health phenotypes. To a lesser extent, it was associated with cognition and BMI. p = non-corrected p-value. * = results unavailable for rs12658032 so a proxy SNP – rs1592757 – was used.

The second of these strongly associated SNPs, rs4593766, is mapped to chromosome 1p31.1 (OR_meta_=1.04(1.03-1.06); P_meta_=4.9×10^−12^) and is similarly located in a region showing no protein-coding annotation and no evidence of brain related eQTL (Table 1; Supplementary File 1: Tables S5-7). The uncharacterised lincRNA transcript *RP4-598G3.1* (A.K.A LINC01360 (ENSG00000233973.6)) is mapped to the region (Table S9), and rs4593766 is in perfect LD with rs6424545, a SNP that impacts the canonical splice site of this lincRNA (Table S11). Based on data from the GTEx Portal, *RP4-598G3.1* is expressed predominantly in the testis with some evidence of expression in thyroid. rs4593766 demonstrates SNP-level associations in mental health related GWAS in addition to ADHD and MDD, such as anxiety and schizophrenia (Supplementary File 3). The region is also associated with years of education, birthweight, BMI and waist circumference.

Nine of the 14 index SNPs were found to be (or be in high LD with) cis-eQTL (see Supplementary File 1: Tables S5-7). Two of these index SNPs (rs2029591 and rs8084351) are linked to brain-eQTL. rs2029591 (OR_meta_=1.05(1.03-1.07); P_meta_=5.9×10^−9^) is mapped to chromosome 3p21.31 and is physically co-located over many genes (Table 1). rs2029591 showed evidence as an eQTL for *AMT, FAM212A, GPX1, NCKIPSD* and *RNF123* across multiple brain tissues (Supplementary File 1: Tables S5-7). As an example, *NCKIPSD* encodes a protein involved in the building and maintenance of dendritic spines and modulates neuronal synaptic activity (33). rs2029591 shows additional SNP-level associations with cognitive traits, BMI and various mental health traits including irritability and bipolar disorder (Supplementary File 3).

rs8084351 (OR_meta_=1.04(1.02-1.05); P_meta_=2.4×10^−9^) is mapped to chromosome 18q21.2 and is physically co-located over the *DCC* gene (Table 1 & Figure 3). *DCC* encodes a receptor involved in guiding neuronal growth and is highly expressed in the brain (34). rs8084351 showed evidence as an eQTL for *DCC* in the cerebellum (b=-0.39, SE=0.08, p=4.60×10^−6^) (Supplementary File 1: Table S5-7). In addition to MDD and ADHD, rs8084351 is also associated at the SNP-level with cognitive traits, neuroticism and mental health traits such as irritability, depressive symptoms and schizophrenia (Figure 4; Supplementary File 3).

**Figure 3.**
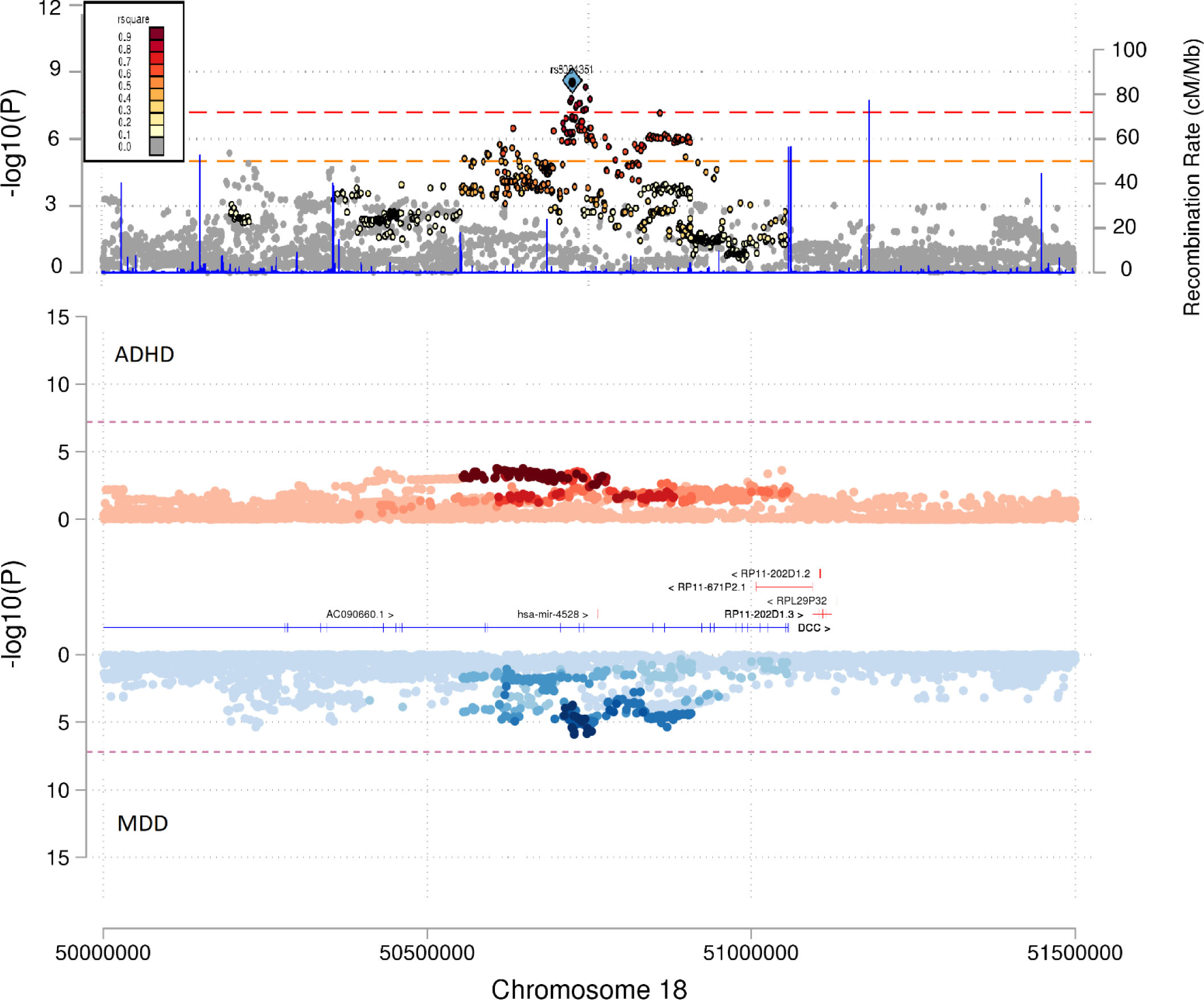
Miami and locus plots showing the association of SNP rs8084351 with ADHD, MDD and the meta-analysis. As shown in the Miami plot, the SNP rs8084351 on chromosome 18 had a subthreshold association with both ADHD (p=4.11×10^−4^) as shown in red and with MDD (p=1.24×10^−6^) as shown in blue. However, it was genome-wide significant in the meta-analysis (p=2.39×10^−9^), as shown in the locus plot. Genes in the surrounding region of 5×10^7^ to 5.15×10^7^ base pairs included *DCC*. Genes are defined according to residence within the LD block of index SNP. Protein-coding genes are shown in blue. Non-protein-coding genes are shown in red. The x-axis is chromosomal position in base pairs and the y-axis is the p value (-log10 p value) of the association of the SNP with both disorders.

**Figure 4.**
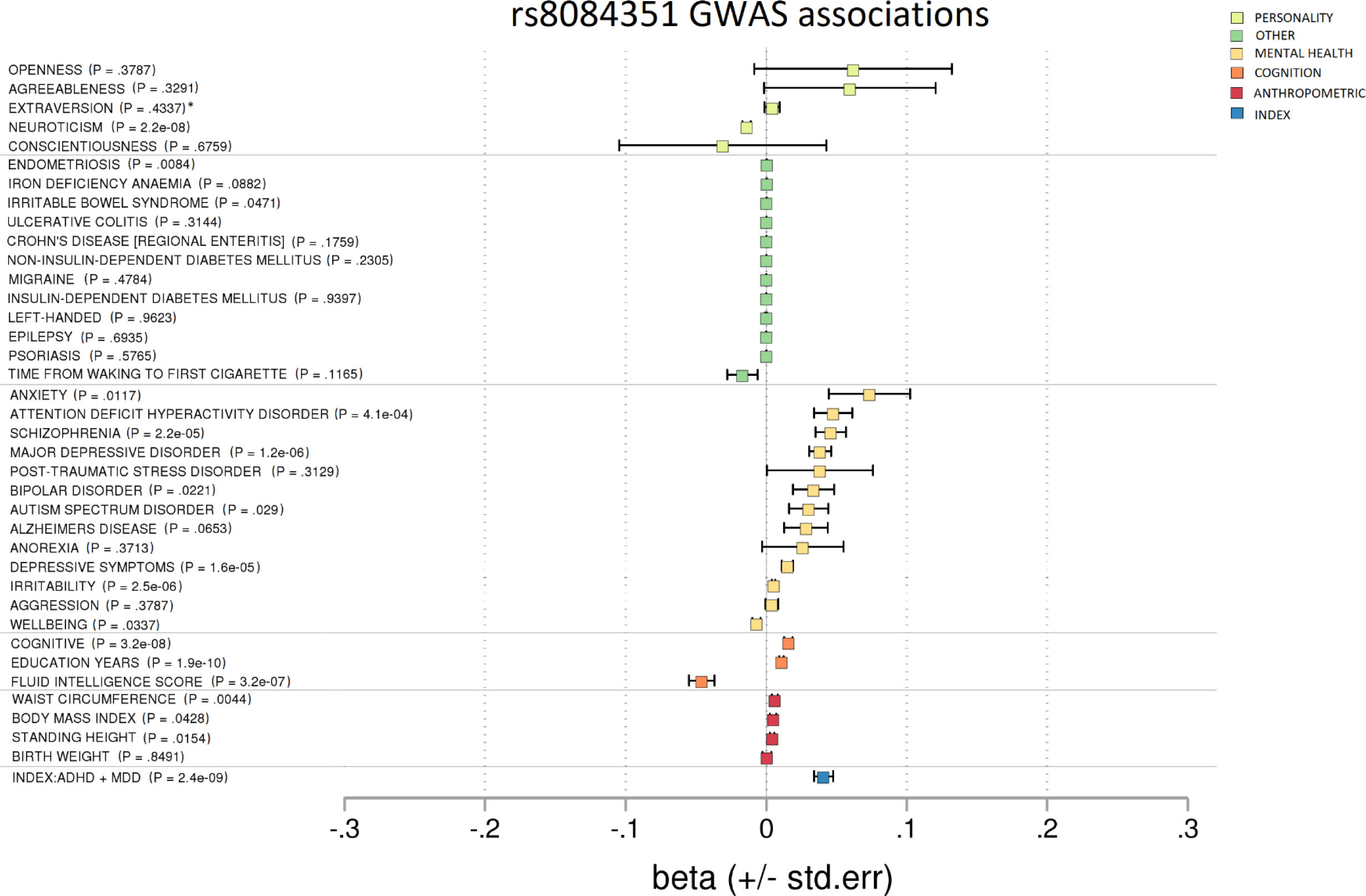
Forest plot showing GWAS associations of rs8084351. To investigate pleiotropy, rs8084351 was tested for associations across 37 phenotypes covering various aspects of health and functioning. rs8084351 was associated mainly with mental health phenotypes. It also demonstrated associations with cognition and neuroticism. p = non-corrected p-value. * = results unavailable for rs8084351 so a proxy SNP – rs7505145 – was used.

Alongside the majority of the SNPs reported in this analysis (i.e. rs61867322, rs2509805, rs113901452, rs1371048, rs8084351, rs139161896, rs2029591, rs2433018, rs12226775, rs71639293, rs16827974 and rs17775184), rs8084351 is an example of a novel finding having been observed below the standard GWS reporting threshold in both the MDD GWAS and ADHD GWAS. These SNPs when combined in this meta-analysis reveal a strong combined signal (see Table 1; e.g. Figure 3).

Detailed description of the annotations for the additional identified markers are given in Supplementary File 2. Meta-analysis results remained similar when z-score thresholds were used to define Model 1, 2 and 3 SNPs instead of p-value thresholds (Supplementary File 1: Table S1).

## Discussion

We performed a GWAS meta-analysis to identify SNPs with a joint contribution to ADHD and MDD. We identified 14 such LD-independent SNPs. All 14 SNPs demonstrated concordant directions of effect for ADHD and MDD. Five of the SNPs were in high LD with nearby SNPs reported as GWS in the individual MDD or ADHD GWAS (13,14) (see Table S6). The remaining 9 SNPs had not been previously reported in either the individual ADHD or MDD GWAS. None of the identified SNPs were genome-wide significant (GWS) in *both* ADHD and MDD GWAS individually.

We examined association across 37 GWAS selected to represent well-powered GWAS of mental health, cognition, personality, anthropometric and other heritable traits to explore the pleiotropy of each identified SNP. In general, for each of the identified SNPs, we observed deviation from the null across traits within the mental health category.

Annotation of GWAS findings can often add supporting data through revealing functionally relevant SNPs, thus moving from associated SNPs to genes and potential biological processes. For the traits under investigation in this study, we annotated the SNP and the associated region (including other SNPs in high LD to the index SNP) for transcripts, eQTL and SNP consequences. Essentially, our approach was to identify SNPs showing evidence of functional impact in trait relevant tissues (i.e. brain) or showing a relationship to genes related to brain function and/or neurodevelopment. Our top finding (rs12658032) revealed none of these properties. The only transcripts in the region (*RP11-6N13.1* and *RP11-6N13.4*) do not show evidence of brain tissue expression. The region that includes rs12658032 is associated with multiple traits and therefore warrants further interrogation. A recent study reported a SNP in this region (rs1363105, a SNP in LD with rs12658032) that was associated with ADHD, MDD, ASD and anorexia (35).

Nine of the observed associations were novel in that they had not been reported previously in the individual ADHD or MDD GWAS. For instance, rs8084351 on chromosome 18 shows a modest subthreshold association with ADHD and a subthreshold association with MDD. However, we observed a strong GWS association of rs8084351 in the meta-analysis of ADHD and MDD. This SNP provides an example of the increased power of meta-analysis to detect variants that contribute to multiple traits. It might also suggest that while variants such as rs8084351 do not greatly contribute to the phenotype of ADHD or MDD individually, they are important in the overlap or comorbid traits of these two disorders specifically. rs8084351 was additionally associated in GWAS of other mental health phenotypes, cognition measures and education related phenotypes. Cognition or education could be an example of a shared risk factor or effect of ADHD and MDD that rs8084351 contributes to. This SNP was found to affect expression of *DCC* in cerebellar brain tissue. *DCC* encodes a transmembrane receptor for netrin 1 and guides axonal growth of neurones towards sources of netrin 1 (34). Mutations in *DCC* have been shown to result in disruption of the midline-bridging neuronal commissures of the brain, causing horizontal gaze palsy, scoliosis and intellectual disability (36).

One additional SNP (rs2029591) was found to be an eQTL for brain expression. rs2029591 was shown to be an eQTL for multiple genes across multiple tissues. With respect to brain tissue, rs2029591 is an eQTL for *AMT, FAM212A, GPX1, NCKIPSD* and *RNF123.* Of these five genes, there is literature to support a potential role in brain disorder for *AMT, FAM212A* and *NCKIPSD. AMT* forms part of the enzymatic system responsible for glycine cleavage in mitochondria (37). *AMT* mutations are associated with glycine encephalopathy – a rare condition associated with brain abnormality and learning difficulties among other traits. However, *AMT* showed evidence of expression across many different tissues and was not specific to the brain. *FAM212A* and *NCKIPSD* showed brain-specific expression. Mutation in the genomic region of *FAM212A* has been observed in a child with CNS abnormalities (38). *NCKIPSD* has been evidenced to be involved in the building and maintenance of dendritic spines and modulation of synaptic activity in neurones (33).

The phenomenon whereby we observe common association between two traits is called pleiotropy. There are several types of pleiotropy and mechanisms that give rise to it (39). We identified regions of pleiotropy based on GWAS of ADHD and MDD. However, overlap exists in the symptoms of different mental health disorders and overlapping genetic associations may not be limited to ADHD and MDD. In order to explore the extent of pleiotropy, we screened identified regions against 37 additional GWAS. As expected, we identified regions that show common association across multiple psychiatric disorders. Some of the relationships may be in part due to examining highly correlated traits (e.g. cognition and fluid intelligence scores, depressive symptoms and MDD, or waist circumference and BMI). Some may be subject to correlation through misclassification bias or confounding. Misclassification may arise between related psychiatric presentations. For example, misdiagnosis of bipolar disorder as MDD in the early stages of treatment seeking is common (40). Confounding might arise where a common association with, for example, cognition and ADHD arises only because of ADHD being associated with both the SNP and cognition independently and not through a causal pathway. Models such as vertical pleiotropy offer an alternate explanation through the role of intermediate traits. For example, rs8084351 was associated with numerous mental health phenotypes as well as cognition. Vertical pleiotropy would be where rs8084351 directly influences traits related to ADHD, which influences cognition, which could in turn impact upon MDD risk in a causal cascade, for example.

There were five instances where our associated regions identified from the meta-analysis overlap with regions found to harbour GWS associations in the individual ADHD or MDD GWAS. Although we focus on the SNP with the strongest association in the meta-analysis in addition to strong associations with ADHD and MDD individually, the assumption that the shared association is driven by a common SNP could be a potential limitation in our interpretation. Specifically, the SNP with the strongest association may not represent the same causal variant being shared between ADHD and MDD. Our associations could potentially represent ‘spurious pleiotropy’, whereby be two different SNPs in the ADHD and MDD GWAS are in high LD with one another (39). Additionally, the MDD GWAS used in this study has a much larger sample size than the ADHD GWAS used. This statistical power imbalance may potentially bias meta-analysis results in favour of the better-powered discovery GWAS (MDD). Indeed, for 11 of 14 reported associations we found lower p-values for MDD than ADHD. To reduce over interpretation of GWS SNPs due to stochastic processes driven by the better statistically powered GWAS, we limited our interpretation to SNPs whereby a lower but modest minimum association (P<5×10^−4^) was observed in both studies individually. There are concerns in the literature regarding reliance on p-values (32) and thus we repeated our meta-analysis using z-scores to define Model 1, 2 and 3 SNPs. Results were very similar, with the z-score approach detecting 13 Model 3 SNPs – all of the 14 SNPs found using the p-value approach except the weakest association (rs17775184).

One potential point of bias is sample overlap. The ADHD and MDD GWAS data were generated from multiple cohorts. It is possible that the contributing cohorts share individuals. We are aware of some overlap in control individuals between constituent cohorts in the ADHD and MDD GWAS studies (Supplementary File 5). However, there is no evidence to support overlap of affected individuals in these studies (personal communication; PGC Data Access Committee). Additional issues to consider are the choice and availability of GWAS data. Some of the GWAS are primarily based on clinical diagnosis (13,14) whereas others used data sourced solely from the UK Biobank (41). Classification and diagnosis in the UK Biobank are often based on self-report. Although not necessarily a limitation, it is important to note that the 23andMe cohort of the MDD GWAS was not included in this study. With this cohort removed, MDD heritability (h^2^=0.15 (0.01)) was higher than reported in the original GWAS (h^2^=0.09 (0.004); (13)). The genetic correlation between ADHD and MDD was also higher (r_g_=0.52) relative to the genetic correlation reported when 23andMe is included (r_g_=0.42) (13,14). One explanation is that the self-reported phenotype used by 23andMe affords a broader spectrum of clinical severity and increases heterogeneity, in comparison to clinician derived diagnosis used by contributors to the non-23andMe MDD GWAS cohorts (13). Finally, it is important to note that the ADHD and MDD GWAS summary data used here were of a European ancestry only, restricting the generalisability of our findings.

Advantages of this study include the use of the most recent and well powered GWAS of MDD (13) and ADHD (14) in an adapted meta-analysis, which provides a novel approach for investigating SNPs that make a joint contribution to the two disorders, compared to previous cross disorder GWAS of numerous psychiatric disorders (e.g. Lee et al., 2019 (35)). The exploration of multiple phenotypes in the study by Lee et al (2019) requires evidence across multiple GWAS and thus might omit regions only impacting ADHD and MDD. The screening of other GWAS associations of index SNPs and regions, instead of standard literature review approaches, allowed investigation free from reporting-bias of the associations of the SNPs identified in the meta-analysis with a broad range of phenotypes.

### Future research directions

Future research directions include the use of in-depth functional analysis of the genomic regions identified here to further explore their biological mechanisms in the overlap of ADHD and MDD. Additionally, the use of Mendelian Randomisation-based techniques to infer whether or not identified variants are causal in ADHD, MDD and/or their overlap would be a useful next step.

## Conclusions

In conclusion, this study highlights 14 LD-independent SNPs contributing to the genetic overlap of ADHD and MDD, 9 of which are novel and did not meet GWS reporting thresholds in either the ADHD or MDD GWAS alone. Compared to existing cross disorder papers, this paper focuses specifically on ADHD and MDD using a method to highlight shared SNPs regardless of direction of effect. The results build upon existing evidence of the genetic architecture of ADHD and MDD, while revealing the regions of the genome that potentially explain some of the overlap observed between the disorders. Findings suggest that there might be some unique genetic architecture to the overlap of ADHD and MDD. eQTL results support the biological relevance of certain identified SNPs to mental health phenotypes. These SNPs seem to be largely specific to mental health related phenotypes rather than other trait classes, supporting the existence of common genetic pathways amongst psychiatric disorders. A greater understanding of the divergence and commonalities of genetic contributions to psychiatric disorders may better inform current diagnostic boundaries.

## Acknowledgements and Funding

This study was supported by a Medical Research Council Centre Grant (No. MR/L010305/1). This publication is the work of the authors and Victoria Powell will serve as guarantor for the contents of this paper.

