## Supplementary File 2 for "Investigating regions of shared genetic variation in attention deficit/hyperactivity disorder and major depressive disorder: A GWAS meta-analysis"

**Annotation of association signals within identified regions method**

#### Gene transcripts

Transcript annotations were assigned based on overlap with the LD-ranges (with a border of 35kb 5’ and 5kb 3’ to the transcript boundaries) using the Homo sapiens GRCh37.87 gene sets (downloaded from <ftp://ftp.ensembl.org/pub/grch37/release-90/gtf/homo_sapiens/Homo_sapiens.GRCh37.87.gtf.gz>).

#### eQTL

Cis-eQTL (expression quantitative trait loci) annotations were assigned to index SNP and SNPs in high LD. All cis-eQTL with a tissue specific P< 1 x 10^-5^ were mapped. Where more than one SNP linked to the index SNP contained an eQTL for a tissue, the strongest association was reported. The eQTL data was derived from the 48 tissue types available on GTeX version 7 (downloaded from <https://www.gtexportal.org/home/>; (32)).

#### Variant Consequence

Variant consequence annotations were assigned to index SNP and SNPs in high LD. Annotations included Consequence, CADD Phred Score, PolyPhen Category and SIFT Category. Consequence report were limited to CONSSCORE greater than 2 including 5PRIME_UTR[3]; REGULATORY[4]; NONCODING_CHANGE, SPLICE_SITE, SYNONYMOUS, UNKNOWN[5]; CANONICAL_SPLICE[6]; FRAME_SHIFT, INFRAME, NON_SYNONYMOUS, STOP_LOST[7]; and STOP_GAINED[8]. All annotations were extracted and mapped from the CADD v1.3 raw data (downloaded from <https://cadd.gs.washington.edu>).

*Chromatin State*

Chromatin state was defined using the ChromHMM annotations (http://compbio.mit.edu/ChromHMM/). ChromHMM integrates multiple ChIP-seq data of various histone modifications to annotate the proportion of putatively active or senescent chromosome regions. Data were extracted and mapped from the CADD v1.3 raw data (downloaded from https://cadd.gs.washington.edu).

**Protein-coding genes within LD range of model 3 SNPs**

**Chromosome 3 – 49193081 to 49890967 bp**

**AMIGO3.** Encodes a member of a family of interacting transmembrane proteins located on the cell surface. This family’s predicted function is in cell adhesion.

**AMT.** Encodes one of four proteins of the glycine cleavage system within mitochondria. Mutations in these proteins have been associated with cases of glycine encephalopathy (GCE). Ubiquitously expressed.

**APEH.** Encodes the enzyme acylpeptide hydrolase, which catalyses terminal acetylated amino acid hydrolysis from small acetylated peptides. Deletions at this locus are associated with decreased activity of this enzyme. It can be important in destroying proteins damaged by oxidation in cells. Deletions of APEH are found in a variety of cancers. Ubiquitously expressed.

**BSN.** Encodes part of a network of pre-synaptic proteins involved in events at the nerve terminal. It is thought to be a scaffolding protein that contributes to the organisation of the pre-synaptic cytoskeleton. Expressed primarily in neurones of the brain.

**C3orf62.** Ubiquitously expressed.

**C3orf84.** Restricted expression in testes.

**CCDC36.** Biased expression in testes.

**CCDC71.** Ubiquitously expressed.

**CDHR4.** Biased expression in lung and testes.

**DAG1.** Encodes dystroglycan – part of the dystrophin-glycoprotein complex linking the extracellular matrix and the cytoskeleton in skeletal muscle. Certain DAG1 mutations are known to cause forms of muscular dystrophy. Ubiquitously expressed.

**FAM212A.** Broadly expressed.

**GMPPB.** Encodes a subunit of the enzyme GDP-mannose pyrophosphorylase, which catalyses the conversion of mannose-1-phosphate and GTP to inorganic diphosphate and GDP-mannose. GDP-mannose is required in glycosylation pathways. Mutations in GMPPB have been identified in patients with muscular dystrophy, in some cases with additional brain abnormalities or mental retardation. Ubiquitously expressed.

**GPX1.** Ubiquitously expressed. Encodes member of glutathione peroxidase family. These enzymes catalyse the reduction of organic hydroperoxides and hydrogen peroxide by glutathione, thus protecting cells from oxidative damage. Studies have indicated that hydrogen peroxide is also involved in other processes including growth-factor mediated signal transduction and mitochondrial function and thus glutathione peroxidases could also affect these functions. Ubiquitously expressed.

**IP6K1.** Encodes inositol triphosphate – a messenger molecule that releases calcium from intracellular stores. Ubiquitously expressed but more so in the brain than other tissues.

**KLHDC8B.** Encodes a protein with a beta-propeller structure of kelch domains, which allows protein-protein interactions to take place. Mutations in this gene have been found in patients with Hodgkin lymphoma. Ubiquitously expressed.

**MST1.** Encodes a protein with a structure similar to hepatic growth factor. The receptor for this protein is RON tyrosine kinase, which stimulates ciliary motility in ciliated epithelial cells of the lung when activated. Biased expression in liver. Ubiquitously expressed.

**NICN1.** Encodes a protein which localises to the nucleus. Ubiquitously expressed but more so in the brain than other tissues.

**RHOA.** Encodes a member of the Rho family of small GTP-ases. These cycle between active and inactive states and function as molecular switches in signal transduction cascades. They are also involved in reorganising the actin cytoskeleton for cell morphology and motility. Overexpression of this gene is associated with proliferation of cancerous cells and metastasis. Ubiquitously expressed.

**RNF123.** Encodes an E3 ubiquitin ligase that functions in the progression of the cell cycle. Ubiquitously expressed.

**RP11-3B7.1.** Encodes a component of the spliceosome complex. It is one of the retinitis pigmentosa-causing genes. Ubiquitously expressed.

**TCTA.** Ubiquitously expressed.

**TRAIP.** Encodes a protein containing an N-terminal ring finger domain. Interacts with TNRF-associated factors (TRAFs), leading to cell apoptosis via nuclear factor kappa-B activation. Mutations in TRAIP have been found in patients presenting with intrauterine growth defects, dwarfism, microcephaly and mental retardation.

**UBA7.** Encodes a member of the E1 ubiquitin-activating enzyme family. It is a retinoid target that triggers cell degradation and apoptosis in acute promyelocytic leukemia. Ubiquitously expressed.

**USP4.** Encodes a protease that deubiquitnates target proteins, maintaining the operations of the endoplasmic reticulum. Ubiquitously expressed.

**NCKIPSD** **(3:48,663,813).** This gene was not located in the LD window but was found to have its expression affected by rs2029591 in eQTL analysis. Encodes a protein with a nuclear localisation signal. It is involved in signal transduction and may be involved in sarcomere maintenance. It is involved in the building and maintenance of dendritic spines and modulates synaptic activity in neurones. Ubiquitously expressed.

**Chromosome 5 – 44816452 to 44944054 bp**

**MRPS30.** Encodes the 28S subunit protein of a mitochondrial ribosome. Mitochondrial ribosomes help in protein synthesis in mitochondria. Ubiquitously expressed.

**Chromosome 5 – 92995013 to 92995013 bp**

**FAM172A.** Ubiquitously expressed.

**Chromosome 10 – 106544216 to 106830537 bp**

**SORCS3.** Encodes a type-1 transmembrane receptor which is a member of the vacuolar protein sorting 10 receptor family. Mutations in this gene are thought to contribute to shared risk across psychiatric disorders. Highly expressed in the brain.

**Chromosome 11 – 48234357 to 49873791 bp**

**FOLH1.** Encodes a type II transmembrane glycoprotein which acts as a glutamate carboxypeptidase on various substrates. It is expressed in a number of tissues but expression in the brain may be involved in pathological conditions associated with glutamate excitotoxicity. An example is motor neurone death in amyotrophic lateral sclerosis. Biased expression in small intestine, prostate and brain amongst others.

**OR4A47, OR4B1, OR4C3, OR4C5, OR4S1, OR4X1, OR4X2.** Each encodes a member of the olfactory receptor family. Olfactory receptors detect odours in the nose to trigger a neuronal response as part of smell perception. Broadly expressed.

**TRIM49B.** Low observed expression biased to brain and testis.

**TRIM64C.** Low observed expression biased to placenta and testis.

**Chromosome 12 – 89721105 to 89904596 bp**

**DUSP6.** Encodes a member of a class of proteins that dephosphorylate MAPK (mitogen-activated protein kinase). Activation of MAPK cascades plays a role in various cellular process including proliferation and apoptosis. Ubiquitously expressed.

**POC1B.** Encodes a protein that localises to the centrioles and has an apparent role in centriole duplication and maintenance. Mutations in POC1B result in cone-rod dystrophy. Ubiquitously expressed.

**Chromosome 15 – 47659445 to 47685378 bp**

**SEMA6D.** Encodes a class 6 transmembrane semaphorin. Semaphorins have been implicated as having roles in axon pathfinding and branching. Transmembrane semaphorins can act as chemorepellants in axon guidance. Broadly expressed.

**Chromosome 18 – 50713243 to 50746748 bp**

**DCC.** Encodes a netrin 1 receptor and guides neuronal axon growth cones towards sources of netrin 1. Mutations of DCC have been found in patients with agenesis of the corpus callosum and patients with gaze palsy, scoliosis and intellectual disability. Biased expression in the testis, brain and lung and adrenal gland.

**A note on sample overlap**

Due to the collaborative nature of large-scale meta-analytic GWAS, sample overlap can occur which may bias results. Of the 59,851 cases in the MDD GWAS excluding 23andMe (13), we believe there is no overlap with ADHD GWAS (14) cases. Of the 113,154 MDD controls, we believe there is potentially 19% overlap with ADHD controls which could create control effects in this study. 17,841 of the MDD controls were from iPSYCH, meaning that up to 15% of MDD controls could overlap with ADHD iPSYCH controls. However, we do not have access to the numbers of controls overlapping between ADHD and MDD GWAS for iPSYCH. For PCG29 samples, there appears to be nominal overlap of 4% of MDD controls with ADHD controls (personal communication; PGC Data Access Committee).
